# Delimitation of the Tick-Borne Flaviviruses. Resolving the Tick-Borne Encephalitis and Louping-Ill Virus Paraphyletic Taxa

**DOI:** 10.1101/2021.06.11.448023

**Authors:** Artem N. Bondaryuk, Evgeny I. Andaev, Yurij P. Dzhioev, Vladimir I. Zlobin, Sergey E. Tkachev, Yurij S. Bukin

## Abstract

The tick-borne flavivirus (TBFV) group contains at least 12 members where five of them are important pathogens of humans inducing diseases with varying severity (from mild fever forms to acute encephalitis). The taxonomy structure of TBFV is not fully clarified at present. In particular, there is a number of paraphyletic issues of tick-borne encephalitis virus (TBEV) and louping-ill virus (LIV). In this study, we aimed to apply different bioinformatic approaches to analyze all available complete genome amino acid sequences to delineate TBFV members at the species level. Results showed that the European subtype of TBEV (TBEV-E) is a distinct species unit. LIV, in turn, should be separated into two species. Additional analysis of the diversity of TBEV and LIV antigenic determinants also demonstrate that TBEV-E and LIV are significantly different from other TBEV subtypes. The analysis of available literature provided data on other virus phenotypic particularities that supported our hypothesis. So, within the TBEV+LIV paraphyletic group, we offer to assign four species to get a more accurate understanding of the TBFV interspecies structure according to the modern monophyletic conception.

## 1. Introduction

Genus *Flavivirus* includes 53 species (as of November 2020) and more than 40 of them are pathogenic for humans. In accordance with a vector, flaviviruses can be divided into the tick-borne flavivirus (TBFV) group, the mosquito-borne flavivirus group, and the no known vector group (Grard et al., 2007; Moureau et al., 2015).

TBFV are a large group of arboviruses transmitted by hard and soft ticks. Members of the TBFV are widely dispersed across Africa, Europe, Asia, Oceania, and North America (Heinze et al., 2012). TBFV may infect vertebrates which can be reservoirs and play a vital role in maintenance of viruses in natural foci.

The TBFV group has 12 members: Louping ill virus (LIV), Kyasanur Forest diseases virus (KFDV), Powassan virus (POWV), Omsk haemorrhagic fever (OHFV), tick-borne encephalitis virus (TBEV), Gadgets Gully virus (GGYV), Langat virus (LGTV), Royal Farm virus (RFV), Meaban virus (MEAV), Saumarez Reef virus (SREV), Tyuleniy virus (TYUV), Kadam virus (KADV); the first five of them (LIV, KFDV, POWV, OHFV, and TBEV) are important pathogens of humans also known as the “tick-borne encephalitis (TBE) serocomplex”: OHFV and KFDV cause haemorrhagic fever in humans, other three viruses (LIV, POWV, and TBEV) induce meningitis, encephalitis, and meningoencephalitis (Shi et al., 2018). The most notorious member of this complex is TBEV. About 12,000 tick-borne encephalitis (TBE) cases are detected annually. Foci of TBEV have been identified in the Russia, Europe, northern China, South Korea, and Japan (Dobler et al., 2017). Recently, Fares et al. (2020) have reported the presence of TBEV (European subtype) in northern Africa (Tunisia).

Not so long ago, taxonomy rearrangements within the TBFV group have taken place. Based on genetic analysis of a polyprotein and an envelope protein of KFDV and Alkhumra haemorrhagic fever virus (AHFV), species *Kyasanur Forest diseases virus* and *Alkhumra haemorrhagic fever virus* have been fused into the one taxon – *Kyasanur Forest diseases virus* (Charrel et al., 2001). Also, considering genetic distances, species *Powassan virus* and *Deer tick virus* have been merged into one as well (Beasley et al., 2001).

Interesting taxonomy perturbations have occurred with species *Royal farm virus* and *Karshi virus*: according to the International Committee on Taxonomy of Viruses (ICTV) these two species had been merged in 1999 with no specific reasons mentioned in the available literature. Here, it is important to note that the phylogenetic distance between RFV and Karshi virus (KFV) significantly exceeds empirical interspecies threshold regarding the other TBFV species (Grard et al., 2007; Moureau et al., 2015).

Another vague situation in terms of taxonomy is observed within the TBEV group: on the phylogenetic trees, LIV is a sister group of the European subtype of the TBEV clade (Dai et al., 2018; Uzcátegui et al., 2012), thus the species *Tick-borne encephalitis virus* is a paraphyletic group. This fact contradicts not only modern cladistics, but also the ICTV definition of the species taxon: “A species is a **monophyletic group** of viruses whose properties can be distinguished from those of other species by multiple criteria https://talk.ictvonline.org/information/w/ictv-information/383/ictv-code”. Therefore, the taxonomy status of the TBEV+LIV group remains unclear. Also, there are several LIV-like viruses (Spanish goat encephalitis virus (SGEV), Spanish sheep encephalitis virus (SSEV), Turkish sheep encephalitis virus (TSEV), and Greek goat encephalitis virus (GGEV)) that are not currently classified.

The intraspecies structure of TBEV is presented by five main subtypes (listed in the order they were discovered and described): the Far-Eastern (TBEV-FE), the European (TBEV-E), the Siberian (TBEV-S), the Baikalian (TBEV-B; Adelshin et al. (2019); Kovalev and Mukhacheva (2017); Kozlova et al. (2018)), the Himalayan (TBEV-H; Dai et al. (2018)). The subtype names point out their prevalent geographic distribution, however, for TBEV-E and TBEV-S, there are “irregular” isolates found far from their primary foci. TBEV-S, in particular, is most widespread, it is found almost across TBEV distribution range. On a phylogenetic tree, the TBEV subtypes are all monophyletic groups and divided by internal branches with the lengths possibly being long enough to delineate these subtypes as species taxa.

In addition to monophyly and genomes relatedness, ICTV also considers the follow criteria: natural and experimental host range, cell and tissue tropism, pathogenicity, vector specificity, and antigenicity.

The final solution on the TBFV taxonomy issue is important concerning epidemiology and prevention. A virus species due to the natural selection obtains specific biological properties allowing them to adopt to specific host range. In the case of TBFV, during infection, primary cell barrier is overcome due to physicochemical interactions between virus envelope glycoprotein (E protein) and receptors on the host cell surface. Amino acid sequences of E protein of different TBFVs determine their specific host range. In humans, E protein is the main target of immune response both after natural infection and vaccination. Several studies showed significant variation of the E protein of TBEV subtypes that reduce cross-immune response to infection by different TBEV strains (Bukin et al., 2017; Rey et al., 1995). Clarification of taxonomy status of different TBFV members can aid universal multivalent vaccine developers to improve prevention of virus infections.

This study aimed to clarify the ambiguity in the taxonomy structure of the TBFV group using three molecular species delimitation methods and all available complete genome data. Then, we focused on the analysis of the most dangerous and widespread group of TBEV (including LIV). For TBEV and LIV we carried out analysis of antigenic determinants of envelope protein (E) to clarify the issue of interspecies position of them. In conclusion, we analysed available literature on the remaining species criteria to make our approach in determining the interspecific threshold more comprehensive and holistic.

## 2. Materials and methods

### 2.1. Genome data set preparation

To delimit species units within the TBFV group, amino acid sequences of a complete ORF (3414 aa) available in ViPR (Pickett et al., 2012) were used. For each species, at least one sequence was found. A total of 278 amino acid sequences were used in the analysis (Table 1).

**Table 1.** The number of amino acid sequences of an ORF region (∼ 3414 aa) for each TBFV group member used for phylogenetic reconstruction and species delimitation.

| Member | A number of ORF sequences |
| --- | --- |
| GGYV | 2 |
| KADV | 1 |
| KFDV+AHFV | 25 |
| LGTV | 3 |
| LIV+LIV-like | 30 |
| MEAV | 1 |
| OHFV | 3 |
| POWV+DTV | 23 |
| RFV+KSIV | 5 |
| SREV | 1 |
| TBEV | 181 |
| TYUV | 3 |
| **Total** | **278** |

Sequences were visualised with AliView v.1.26 (Larsson, 2014) and aligned with MAFFT v.7 online (Katoh et al., 2017; Kuraku et al., 2013).

### 2.2. Phylogenetic analysis and model selection

Phylogenetic analysis was performed with BEAST v.1.10.4 (Suchard et al., 2018) and IQTREE v. 1.6.12 (Nguyen et al., 2015) as implemented on the CIPRES web server (Miller et al., 2010). The best-fit amino acid substitution matrix with the lowest value of the Bayesian information criterion (BIC) (FLU+G_4_+I) was chosen by ModelFinder implemented (Kalyaanamoorthy et al., 2017); number of gamma categories, alpha (shape) parameter of a gamma distribution (0.78) and proportion of invariant sites (0.13) were fixed in further analysis in BEAST. Based on coefficient of variation values of substitution rates (a mean value = 0.58; 95% HPD, 0.49-0.68), the relaxed clock with an uncorrelated lognormal distribution (UCLD) was selected as a molecular clock model. The birth-death (BD) model was chosen over a Yule prior since a preliminary BEAST run has demonstrated that a 95 % highest posterior density (HPD) interval of death rate lay far enough from zero (0.991-0.999).

The reproducibility of each Markov chain Monte Carlo (MCMC) analysis was tested by five independent BEAST runs. Each MCMC analyses was run for 100 million iterations, with a tree sampled every 2,500 steps. Burn-in proportion was selected in each run individually. Then, we combined all analysis logs (*.log and *.trees file of the BEAST output) in LogCombiner and analysed the summary log with TreeAnnotator to obtain the most credible clade tree. The convergence and effective sample sizes (ESS) of the summary log were assessed using a Tracer v.1.7.1 program (Rambaut et al., 2018). The BEAST project file, the consensus tree and the output Tracer logs (combined by LogCombiner) are available from https://doi.org/10.6084/m9.figshare.13614080.

### 2.3. Species delimitation

To delineate TBFV species we employed three bioinformatics delimitation methods. The maximum likelihood tree, reconstructed in IQTREE, was rooted by Apoi virus (NC_003676) as an outgroup and used to delimit species by a Bayesian implementation of the Poisson tree processes (PTP) model (Zhang et al., 2013) using an online service: https://species.h-its.org/.

The generalized mixed Yule coalescent (GMYC) method (Fujisawa and Barraclough, 2013) implemented in the “splits” package for the R was applied to determine clusters at the species level on the ultrametric tree previously reconstructed with BEAST.

The amino acid distance matrix calculated by the maximum likelihood method implemented in IQTREE program was used for species delimitation by using the Automatic barcode gap discovery (ABGD) method (Puillandre et al., 2012) with the online service: https://www.abi.snv.jussieu.fr/public/abgd/.

### 2.4. Comparative analysis of viral antigenic determinants sequences

The TBEV Sofjin strain (1488 nt) was used for a nucleotide BLAST (http://blast.ncbi.nlm.nih.gov/Blast.cgi) to search for homologous E gene nucleotide sequences in GenBank (http://www.ncbi.nlm.nih.gov/genbank/). BLAST parameters were set as following: word size was set as 11; match/mismatch scores – 2,-3; gap costs – existence: 5, extensions: 2. Initially, 982 nucleotide sequences were found. The nucleotide data set obtained was translated to amino acids and filtered by a length threshold of 367 aa (∼75 % of E protein). Using the resulting data set (932 sequences), we performed phylogenetic analysis with IQTREE v.1.6.12 program to determine a virus subtype. As a result, we assigned the next five phylogenetic groups: TBEV-FE, TBEV-S, TBEV-E, TBEV-B and LIV. TBEV-H and other TBEV lineages were excluded cause of an insufficient number of sequences for inter- and intragroup genetic analysis. After that, based on published crystallography results (Rey et al., 1995), fragments exposed at the virus surface - the antigenic determinants, - were defined and isolated from full-length amino acid sequences of the E protein (the length of antigenic determinants was 224 aa; for more information on this procedure see Bukin et al. (2017)). For the amino acid sequences of antigenic determinants, a length threshold was set as 190 aa (85% of total determinants length). The final alignment comprised 812 antigenic determinant amino acid sequences of TBEV and LIV (Table 2).

**Table 2.** The number of sequences used in antigenic determinants comparative analysis.

| Phylogenetic group | Number of sequences |
| --- | --- |
| TBEV-FE | 293 |
| TBEV-S | 159 |
| TBEV-E | 308 |
| TBEV-B | 12 |
| LIV | 40 |
| **Total** | **812** |

To calculate the inter- and intragroup pairwise protein evolutionary distances for antigenic determinants, we performed phylogenetic analysis by IQTREE v.1.6.12 with ultrafast bootstrap replicates (1000 trees in total) (Hoang et al., 2018). The best-fit amino acid substitution matrix according to the lowest BIC values calculated by ModelFinder was HIVb+G_4_. Further, inter- and intragroup pairwise distances with 95% credible intervals (CIs) in each tree were analysed with an R script. The R script is available from https://doi.org/10.6084/m9.figshare.14774094.v1. The *F*_*st*_ values (the measure of intergroup subdivision) were obtained as follows:

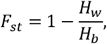

where *H*_*w*_ are mean intragroup genetic distances, *H*_*b*_ are intergroup genetic distances (Hudson et al., 1992). The P-values were determined as a proportion of negative or zero *F*_*st*_ values from the total number of tree samples (1000).

Visualization of violine plots was executed in R with the Vioplot v.0.2. package (https://github.com/TomKellyGenetics/vioplot).

## 3. Results

### 3.1. Phylogenetic analysis

Results of phylogenetic analysis performed in BEAST revealed a clear asymmetric tree shape with very high posterior probability (pp) of main nodes except for the KADV isolate with pp of 0.46; its phylogenetic position regarding the other TBFV members remains uncertain (Fig. 1). All species clusters reviled (which are not singletons) have pp of 1.

**Fig. 1.**
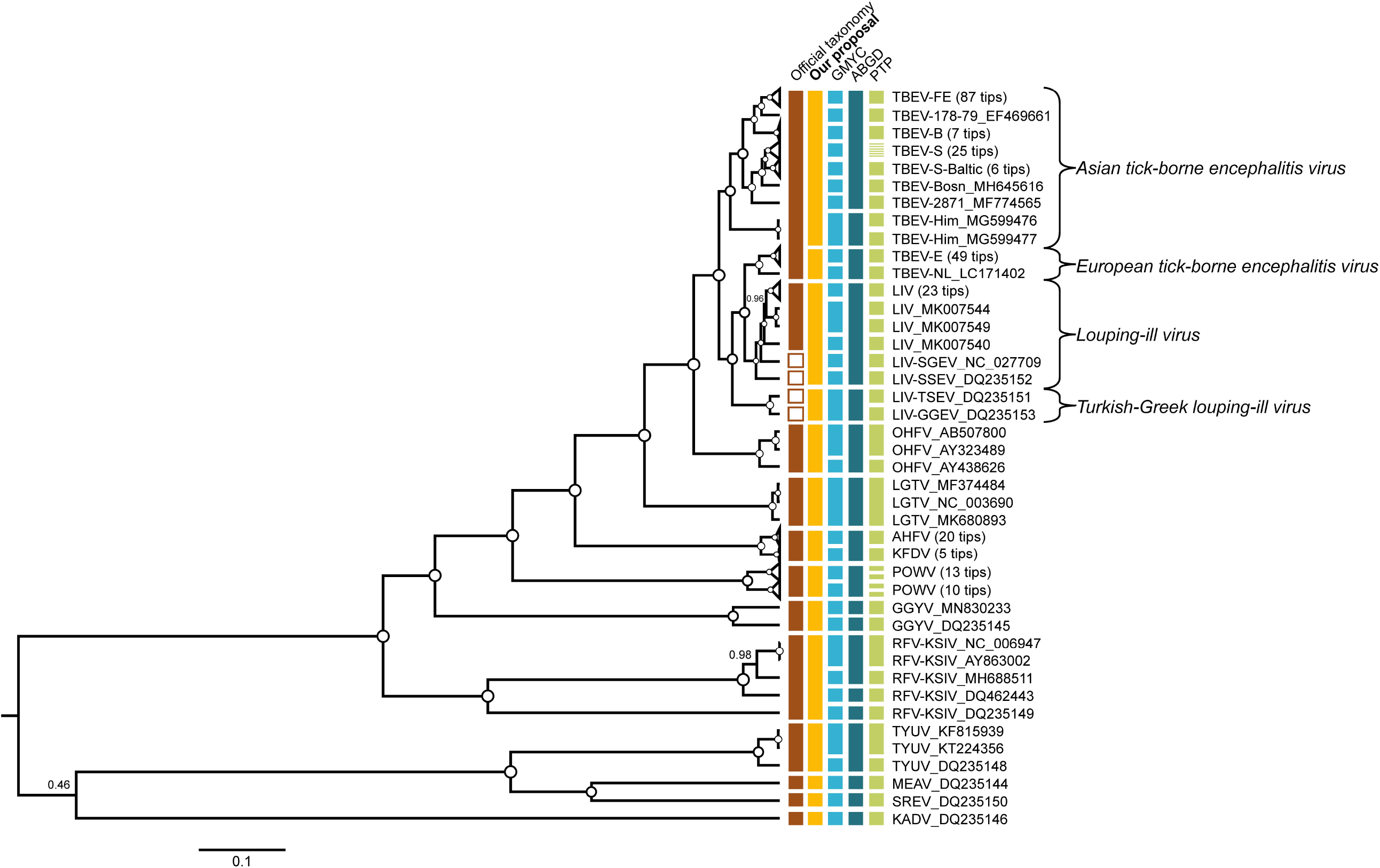
The phylogenetic tree of TBFVs. The tree was reconstructed in BEAST using complete amino acid sequences (n = 278) of the polyprotein (3414 aa). For clarity, some of the wide clades were collapsed. Vertical bars to the right of tree tips indicate official classification (brown), our taxonomy proposal (orange), and delimitation results. Internal nodes with pp = 1 are marked as white circles, otherwise support values are shown by numbers ranged from 0 to 1.

### 3.2. Delimitation results and discrepancies with the official taxonomy

The TBFV group was divided into 34, 18, 44 evolutionary significant species units by the GMYC, ABGD and PTP analysis, respectively (Table 3).

**Table 3.** Results of the species delimitation tests in the TBFV group

| Gene | A number of species clusters |  |  |
| --- | --- | --- | --- |
|  | GMYC (P value for LR-test) | ABGD | PTP |
| Polyprotein | 34 (1.3E-9) | 18 | 44 |

The discrepancy between the official taxonomy and the delineation results is observed within the TBEV+LIV paraphyletic group and OHFV, KFDV+AHFV, POWV, GGYV, RFV+KSIV, TYUV monophyletic groups (Fig. 1).

The TYUV cluster was divided into two species units according to GMYC and PTP methods. The isolate DQ235148 from the Three Arch Rocks National Wildlife Refuge (USA, Oregon) was separated from two other isolates (KF815939, KT224356) from the Russian Far East (the Sea of Okhotsk, Tyuleny Island), with air distance between these isolation places being about 6,000 km.

The RFV+KSIV isolates from the Central Asia (Uzbekistan, Afghanistan, Turkmenistan, and Northwest China) showed significant intragroup amino acid diversity despite their geographic distance is relatively small. Notably that the isolate DQ235149 (Afghanistan) diverged from other RFV+KSIV members more than TUYV diverged from SREV and MEAV. The GMYC, the ABGD and the PTP algorithms split the RFV+KSIV cluster into 4, 3, 4 distinct species units, respectively.

The GGYV cluster formed by two isolates (MN830233, DQ235145) from Australia and Antarctica has split into two species units by all three methods.

The POWV group has split into two species units by the GMYC method, however the PTP algorithm discovered 4 species. The POWV group consists of two distinct clades, which have a clear geographical determinant (the Russian Far-East and USA) of isolates clustering (Supplemental Fig. 1). The PTP method delimits the each of two main cluster into two species units. The GMYC methods identified two main clusters as two species units without splitting them within. ABGD showed the most conservative point of view and didn’t split POWV cluster as it is in the official taxonomy. Thus, delimitation methods remained POWV taxonomy structure unresolved.

According to GMYC and PTP, the KFDV+AHFV cluster is delimited into two distinct species – KFDV (India) and AHFV (Saudi Arabia). In turn, the ABGD support official taxonomy status of the RFDV+AHFV group as a single species taxon.

The OHFV group was divided into two species units by the GMYC and PTP methods, and the ABGD method defined the cluster as a single species.

Delimitation methods showed the most inconsistency in the case of the TBEV+LIV paraphyletic group. All three delineation methods defined TBEV and LIV as distinct species, however interspecies separation in both viruses was different. Generally, within the TBEV group, the GMYC and PTP methods, compared to ABGD, delimited species more frequently – 10, 16, and 3 species, respectively. Concerning LIV and LIV-like viruses, ABGD once again was more conservative (2 species clusters – LIV + SGEV + SSEV and TSEV + GGEV), whereas GMYC and PTP determined 6 and 8 species, respectively. Notably, TSEV isolated in Turkey and GGEV isolated in Greece were not only delimited by all of three methods but they were also geographically distant from LIV (British Isles) and both LIV-like viruses (Spain). In this case, phylogenetic analysis provided strong evidence of geographic clustering, which is also consistent with delimitation analysis. Since the TBEV+LIV group are most representative in terms of a number of available sequences and literature information on the other viral species criteria (e.g., pathogenicity, cell and tissue tropism, vector specificity, etc.), we decided to do the additional analysis of this group. We have analysed the envelope protein amino acid sequences of TBEV and LIV to distinguish these putative viruses considering their antigenic properties.

### 3.3. Comparing antigenic determinants of TBEV and LIV

The analysis of protein evolutionary distances between the antigenic determinants of TBEV and LIV showed that LIV is statistically different from all TBEV subtypes (including TBEV-E; Fig. 2, Table 4).

**Table 4.** Inter- and intragroup pairwise genetic distances of TBEV and LIV antigenic determinants

| Comparing virus pairs | Mean intragroup distance | Mean intergroup distance | F<sub>st</sub> | P-value |
| --- | --- | --- | --- | --- |
| S-B | 0.0062 | 0.0115 | 0.3666 | 0 |
| FE-B | 0.0047 | 0.0096 | 0.3968 |  |
| FE-S | 0.0102 | 0.0203 | 0.4424 |  |
| *E-S | 0.0088 | 0.0327 | **0.7003** |  |
| FE-E | 0.0073 | 0.0410 | **0.7961** |  |
| **E-LIV** | 0.0122 | 0.0817 | **0.8424** |  |
| S-LIV | 0.0152 | 0.1080 | **0.8541** |  |
| FE-LIV | 0.0136 | 0.1163 | **0.8773** |  |
| **E-B** | 0.0033 | 0.0322 | **0.8794** |  |
| **B-LIV** | 0.0096 | 0.1075 | **0.9069** |  |

\* - pairs with TBEV-E or/and LIV were bolded

**Fig. 2.**
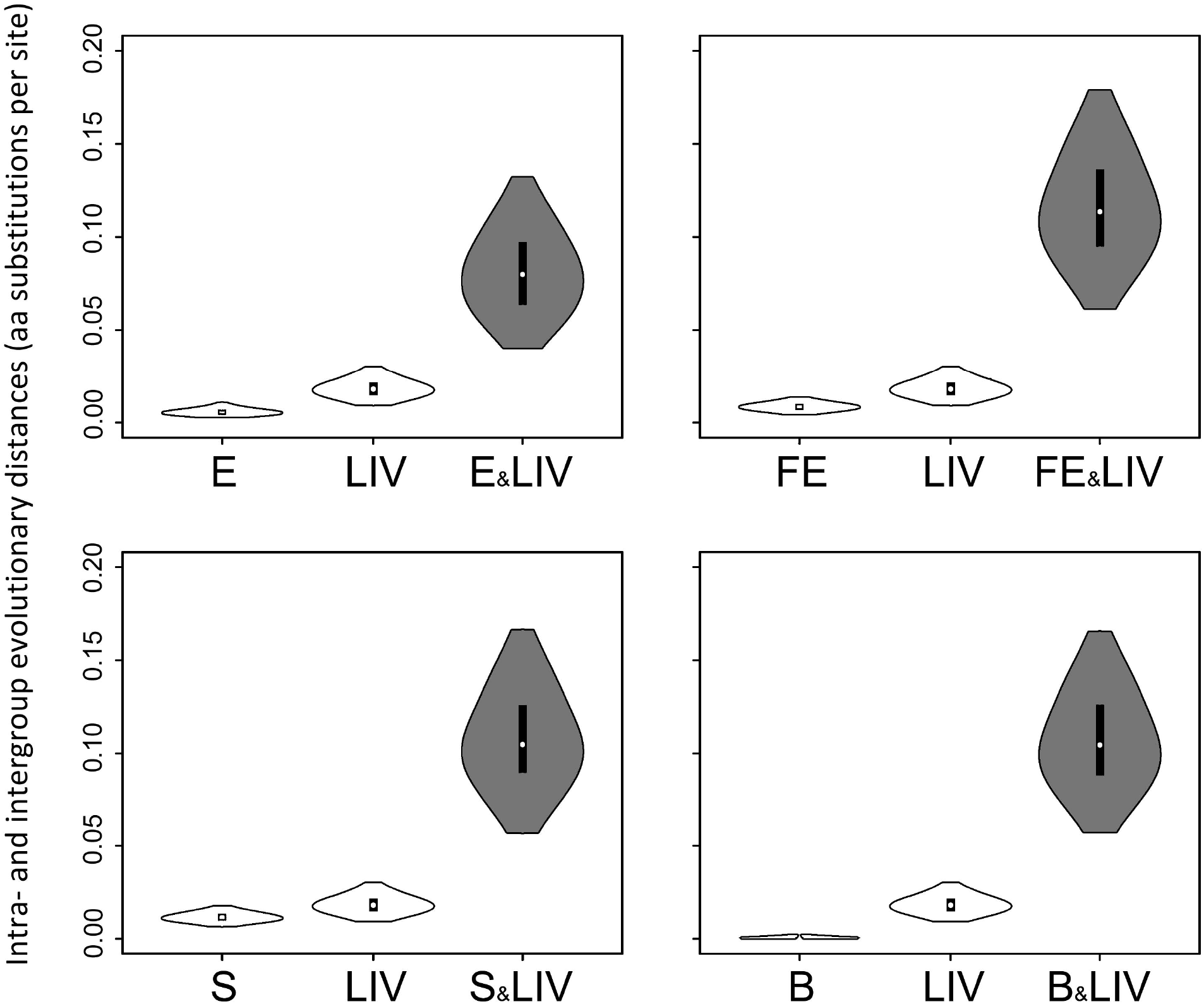
Comparing intra- and intergroup pairwise genetic distances of LIV and TBEV epitopes (224 aa). The distances were calculated based on 1000 replicates of ultrafast bootstrap analysis using 812 amino acid sequences. Distributions of intra- and intergroup pairwise distances are displayed as white and grey violin plots, respectively. Upper and lower boundaries of violin plots represent 95% CIs. Black vertical bars within plots are standard deviation, white circles – a mean value. The Y axes show genetic distances expressed in amino acid residue substitutions per site. On the X axis are TBEV subtypes (FE – Far-Eastern, S – Siberian, E – European, B – Baikalian) and LIV.

LIV intragroup distances have the highest mean value and the widest 95% CI that, in turn, indicate the highest antigenic polymorphism of LIV (Fig. 3). However, 95% CI of LIV are visibly overlapped with TBEV-FE and TBEV-S CIs.

**Fig. 3.**
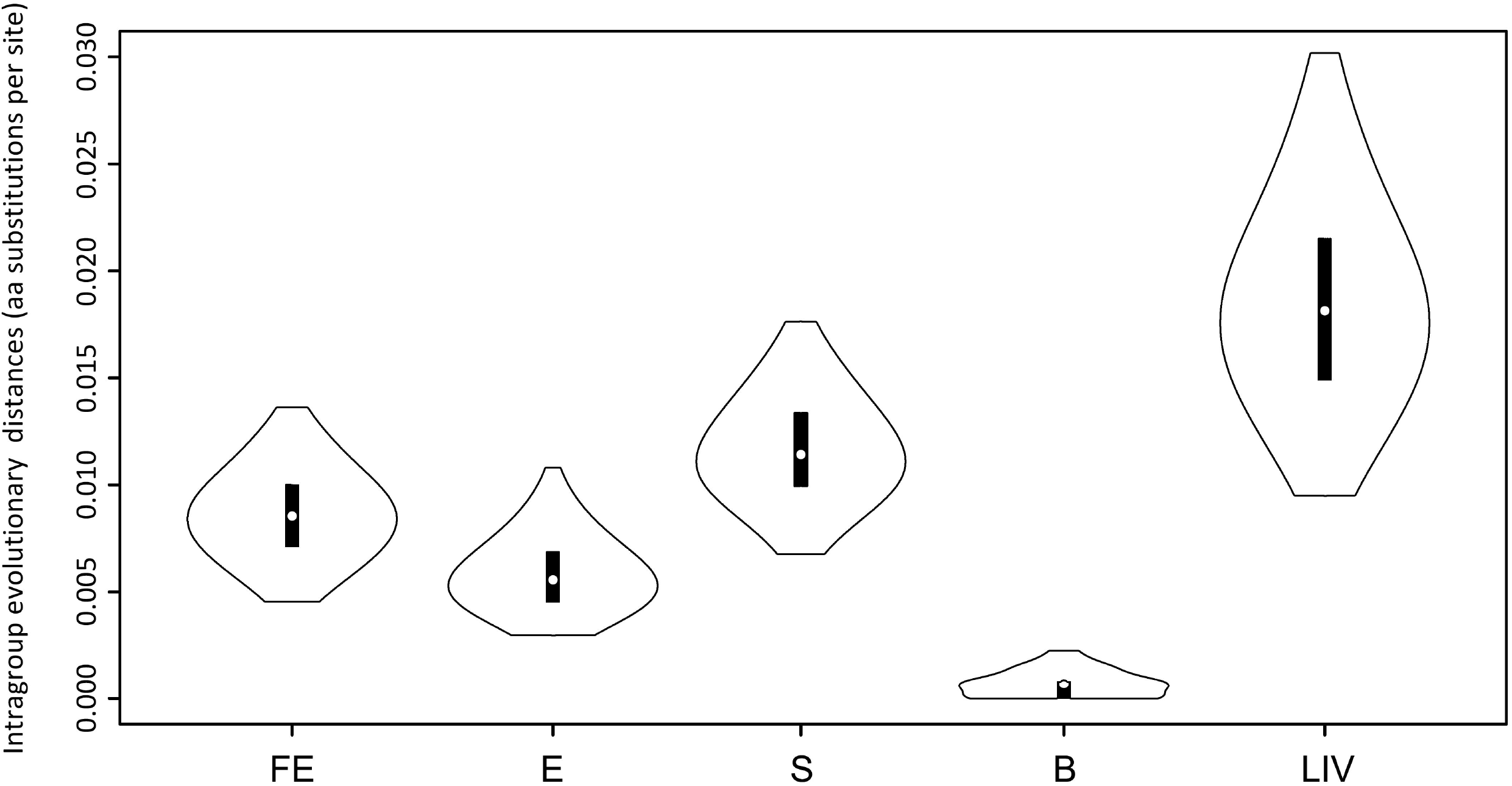
Distributions of intragroup pairwise genetic distances of LIV and TBEV antigenic determinants (224 aa). The designations are the same as in the figure 2.

As with LIV, TBEV-E are significant different from other TBEV subtypes (Supplemental Fig. 2c, g), with an exception of TBEV-S (there is a slight overlapping of 95% CIs; Supplemental Fig. 2e). In contrast, the inter- and intragroup genetic distances of the remaining three TBEV subtypes (TBEV-FE, TBEV-B, TBEV-S) do not significantly differ (there is overlapping of CIs; Supplemental Fig. 2b, d, f).

In all cases of comparing TBEV-E and LIV versus other TBEV subtypes, *F*_*st*_ values were more than 0.5 (Table 4). Notably, in the same table we can see that the mean values of intergroup distances of all viruses analysed are more than intragroup distances, but, as we discussed above, CIs of TBEV-FE, -B, and -S pairwise intragroup distances are overlapped significantly and, as a consequence, are not statistically distinct.

On the consensus phylogenetic tree (Supplemental Fig. 3) of TBEV and LIV reconstructed using the amino acid sequences of antigenic determinants, TBEV-S has no reliable bootstrap support (91 and 45 score for ultrafast bootstrap and SH-aLRT methods, respectively) and, therefore, cannot be surely separated from TBEV-FE and -B unlike TBEV-E and LIV which have high enough support score (95/94 and 99/100, respectively).

## 4. Discussion

### 4.1. Delimitation of TBEV and LIV phylogenetic group

All three delimitation methods separated TBEV-E into a distinct species taxon. The results of the TBEV and LIV antigenic determinants comparison demonstrated that TBEV-E and LIV are probably different from each other and the remaining TBEV subtypes regarding their antigenic properties. Concurrently, TBEV-FE, -B, and -S subtypes are not statistically distinct by antigenic determinants structure. To keep monophyly principle, we should whether to combine TBEV-E and LIV as a single species, or assign them as two separate taxa. Thereby, to holistically scrutinised this problem, we have analysed available literature for other viral species criteria. It should be noted that according to all three delimitation methods TBEV-H was delineated as separated species as well, however, there is no data on its biological particularities and we will therefore not consider it as an independent taxon.

### 4.2. Consideration of the biological and ecological peculiarities of TBEV and LIV and comparing them with species delimitation results

It is reliably known that TBEV infects humans causing severe meningitis, encephalitis, and meningoencephalitis. Contrarily, cases of LIV infections in humans are relatively rare, with signs of acute meningoencephalitis and poliomyelitis being described. The clinical picture for humans infected with LIV is very similar for that produced by TBEV-E: The first phase of disease is characterised by fever (2–11 days) followed by remission (5–6 days) and then the re-emergence of fever and meningoencephalitis lasting 4–10 days, usually with full recovery (Gritsun et al., 2003). Notably, a biphasic course is observed in 74% of TBE patients infected with TBEV-E (Kaiser, 1999), but TBEV-FE and -S infections are predominantly monophasic, only a small reminder demonstrating a biphasic pattern (Mansfield et al., 2009). Also, infections with TBEV-FE often cause an illness with a gradual onset, more severe course, higher rates of severe neurologic sequelae compared to TBEV-E infections (Bogovic and Strle, 2015).

One of the most important peculiarities of neurotropic viruses is their ability to across the blood brain barrier (BBB) and cause encephalitis. LIV induce encephalitis in sheep annually, morbidity and mortality rates ranging from 5 to 60% (Jeffries et al., 2014). In contrast, TBEV seems to show nonvirulence for livestock (there are no reports of mass epizootics in Eurasia) and, apparently, persists in wild rodents asymptomatically. In experiments, it was demonstrated that three TBEV subtypes (TBEV-FE, -E, -S) are able to induce encephalitis in bank voles (*Myodes glareolus*) but causing neuronal death in this natural host in very rare cases (Tonteri et al., 2013). In the same study it was shown that TBEV-FE has distinctive infectious kinetics in bank voles: The long duration of TBEV-FE viremia may suggest a different transmission pattern as compared to TBEV-E.

Intriguing results were obtained during experiments with sheep (Votiakov et al., 2002). In the number of studies, sheep received virus solution containing TBEV-FE and TBEV-E subcutaneously and by intracerebral infection (15 and 24 sheep for TBEV-FE and -E experiments on intracerebral infection, respectively). It was shown that subcutaneous infection of sheep as well as infection via ticks with TBEV-E didn’t yield transition of BBB induced only meningitis. In the case of intracerebral infectious, TBEV-E showed clear biphasic course with neurological symptoms (anisocoria, ptosis, myelitic paresis, tonic-clonic spasm) and mortality rate only 12,5%. The viral titer in the different parts of brain (cortex, cerebellum, medulla, cervical, lumbar) during the first phase (fever phase) ranged 1.2-2.8 lgLD_50_ (mean = 2.0). In the blood, virus titer averaged 2.5 lgLD_50_ and was higher than in the CNS. The morphological studies of the CNS showed glial nodes took 8.5-21.5% of microscope fields of view (FOV; mean – 14,6%). Neuronophagia took only 5.6-10.0% of FOV (mean – 7.4%). Damage to the neurons occurs only in some animals as a secondary inflammatory effect arising from infection of glial cells. Whereas, in contrast, the intracerebral infection of sheep with TBEV-FE (the strain “198”) demonstrated mortality rate of 100%. The course of the disease was monophasic and severe and developed rapidly. Already on days 2-3, signs of focal brain damage simultaneously with fever were rapidly developing in sheep. The viral titer in the CNS was on average 1.3 lgLD_50_ higher (1.2-1.8 lgLD_50_) than in the blood. Primary degenerative CNS disorders prevailed. Neuronophagia took 32.5-42.5% of FOV (mean – 37.5%). Glial nodes ranged widely 34.0-99.0% of FOV (mean – 56.1%). For details, see Supplemental table 2.

Votiakov et al. (1978) also showed that European virus initially did not replicate in or damage neuronal cells even after intracerebral infection. Instead, the primary target of European virus was lymphoid tissue and the virus subsequently appeared in the brains, 6–9 days after inoculation (in cerebellum predominantly) of those animals that developed encephalitis. In turn, Far-Eastern virus directly infected and damaged neurons in the brain, resulting in severe encephalitis. These facts evidently indicate that TBEV-FE is more neurotropic than TBEV-E.

In USSR, in the periods 1954-1961 and 1966-1968 Votiakov V.I. and Protas I.I. conducted clinical observations of TBE in humans in Belarus (BSSR) and Khabarovsk Krai (the Far-East region, where pioneering TBEV research work was done by Lev Zilber’s expedition) (Votiakov et al., 2002). In Belarus, 198 individuals were observed during 1954-1961, and, furthermore, Votiakov et al. continued their observation until 1998 with a number of patients reaching 928. In Khabarovsk Krai (1966-1968), total number of individuals observed was 149. All of them had a contact with ticks. Observations showed clear differences in forms of the disease between cases in Belarus and the Far-East. In Belarus (1954-1961), no cases of encephalitis were reported (meningoencephalitis had a diffuse form with transient signs (32%), whereas focal forms were absent), with serous meningitis prevailing (52%). Furthermore, Votiakov et al. (2002) also reported that for 44 years of observations, no cases of meningoencephalitis with focal forms was registered in Belarus. Opposite, in Khabarovsk Krai (1966-1968), serous meningitis was absent, meningeal form with encephalitis signs predominated (32.2%); together with that diffuse and focal meningoencephalitis were observed (16.1% and 14.1%, respectively). Moreover, in Belarus (1954-1998), no cases of polioencephalomyelitis were observed, unlike Khabarovsk Krai where this form of the disease accounted for 21.5%. For details, see Supplemental table 1. Also, Votiakov et al. (2002) provide data on the number of cases of laboratory-acquired infection in their laboratory in Belarus (9 cases), in laboratories of Slovakia (11 cases), and 10 cases of infections with far-eastern TBEV strains in the Far-East. Five of ten cases involved the far-eastern strains were lethal, while in Slovakia and Belarus no lethal cases were registered. As it was described above, the Far-Eastern strains induced focal meningoencephalitis (4 of 10 cases) and polioencephalomyelitis with wide paresis and bulbar palsy. In Slovakia and Belarus, none of those clinical forms was observed, with meningitis prevailing (6 of 11 cases in Slovakia, 5 of 9 cases in Belarus). The given data once again indicates a more severe nature of the course of TBEV-FE and its relatively high neurotropicity.

Case fatality rates (CFR) for each TBEV subtype are different – TBEV-FE has the highest fatality rate (∼20%) (Bogovic and Strle, 2015; Dobler et al., 2017), TBEV-S CFR rarely exceeds 6-8% (Gritsun et al., 2003), and TBEV-E CFR ranges from 1-2% (Bogovic and Strle, 2015). In turn, only one fatal case of LIV infection has been recorded (Williams and Thorburn, 1962). It should be mentioned that epidemiological data show wide variability – the findings may be biased by the different types of medical treatment and by the distinctions in evaluating abortive and asymptomatic forms proportion of TBE.

The other important particularities of virus species are vector and geographic distribution. It is interesting that for TBEV and LIV ticks are both a vector and a reservoir. It was found that the main vector for TBEV-FE and TBEV-S is *Ixodes persulcatus*, for TBEV-E and LIV, in turn, – *I. ricinus*. The spatial distribution of TBEV and LIV is based mainly on the habitats of these tick species and reservoir hosts: TBEV-E is predominantly distributed in Central Europe, TBEV-S and TBEV-FE are mostly spread throughout Siberia and the Far East. LIV is primarily found in the British Isles (upland areas of Great Britain and Ireland), with records also from the Russian Far-East (Leonova et al., 2015), Norway and Spain (Jeffries et al., 2014). Not so long ago, it was believed that TBEV is absent in the British Isles, though it has been recently shown the presence of TBEV-E in the East of England (Holding et al., 2020).

Considering reservoir transmission hosts, LIV once again demonstrates an obvious difference from TBEV. Unlike all TBEV subtypes, LIV is primarily found in red grouses and sheep inducing encephalitis and high mortality rate in both (78% in red grouses (Gilbert, 2016), 5–60% in sheep (Jeffries et al., 2014)), not small rodents. Although rodents such as field voles (*Microtus agrestis*), bank voles (*M. glareolus*) and wood mice (*Apodemus sylvaticus*) raised an antibody response to infection, they could not produce a substantial viremia and did not support non-viraemic transmission between co-feeding ticks (Gilbert et al., 2000). Furthermore, red grouses tend not to feed adult *I. ricinus* and is not therefore able to maintain transmission cycle without aid of another host that feeds adult ticks (e.g. deer, so-called “reproduction hosts” (Gilbert, 2016)). This leads to the fact that LIV has patchy spatial distribution with different combinations of reservoir hosts occurring. This is exactly opposite of the TBEV transmission patterns and natural foci structure formed by primarily small rodents.

The dissimilarity of the clinical picture, cell tropism, host range specificity, and pathogenicity of TBEV subtypes and LIV may speculatively be explained by differences in the antigenic determinants structure. Hubálek et al. (1995) employed indirect immunofluorescence test and revealed a clear difference between LIV strains and the TBEV-FE prototype strain “Sofjin”. On the UPGMA tree, representing of antigenic relationships of the viruses, LIV strains formed a common cluster with a TBEV-E strain, the TBEV-FE strain laying far from them as an outgroup. It is consistent with our results of TBEV and LIV antigenic determinants comparative analysis (Fig. 2).

The data reviewed supports the hypothesis of considering TBEV-E and LIV as distinct virus species.

### 4.3. Delimitation of the remaining members of the TBFV group

The delimitation methods elucidated cryptic species within following clades: TYUV, RFV+KSIV, GGYV, POWV, KFDV+AHFV, and OHFV. In all of these cases, phylogenetic species concept (members descend from a common ancestor) is kept. Some of the clades (e.g. RFV+KSIV, GGYV) contain distances that obviously exceed the interspecies threshold. In some cases, the situation with cryptic species still remains uncertain.

### 4.4. Our taxonomy proposal

Taking into account all the phenotypic manifestations of viruses described above as well as our analysis results, we offer to delineate TBEV-E (with the NL lineage) and LIV into two distinct taxa from the joint TBEV clad and assign them as *European tick-borne encephalitis virus* and *Louping-ill virus*, respectively. To keep the conception of monophyly, we propose to join SGEV and SSEV with the LIV clade into the single species. As a consequence, to maintain monophyly, TSEV and GGEV should be considered as a distinct species as well (Fig. 1) and assigned as *Turkish-Greek louping-ill virus*. The other TBEV subtypes (TBEV-FE, TBEV-B, TBEV-S, TBEV-H) being a monophyletic group are treated by us as a single species (Fig. 1) which we propose assign as *Asian tick-borne encephalitis virus*.

Consideration of taxonomic status of the other virus species outside the TBEV complex is beyond the scope of this study, however our delimitation results indicate possible fields of future TBFV taxonomy investigation.

## 5. Conclusion

To summaries, we have put the data on all TBEV and LIV particularities observed into a Supplemental table 3. LIV has shown clear differences in severity of the disease in humans and sheep being biphasic like TBEV-E. LIV also has relatively low CFR (only one official recorded case). In our analysis, all three delimitation methods showed that LIV and TBEV-E are distinct species. Comparison of envelope protein amino acid distances elucidated that LIV and TBEV-E are significantly different from all TBEV subtypes (and from each other). TBEV-E as well as LIV has a biphasic course in humans, less disease severity with meningitis prevailing, low CFR, and shares with LIV the common vector – *I. ricinus*. However, experiments with sheep showed that TBEV-E cannot cross BBB, had relatively small lethality rate (even after intracerebral infection), and didn’t cause encephalitis and death after subcutaneous infection or infection via ticks.

We believe that the differences described above are sufficient to delineate TBEV-E and LIV from the joint TBEV clade into distinct taxa.

Considering TBEV-E as a separate species is also of practical importance. In particular, two commercially available vaccines for TBEV prevention based on the K23 and Neudoerfl strains (TBEV-E), namely Encepur (“Novartis Vaccines and Diagnostics” Germany) and FSME Immun Inject (“Baxter”, Austria) have been used in Russia (Siberia region) until 2015 despite that TBEV-E isn’t widespread in this territory (in the literature, there are only isolated cases are described (Adelshin et al., 2015; Demina et al., 2010; Demina et al., 2017)). Our results showed significant differences in phylogenetic pairwise distances of antigenic determinants of main TBEV subtypes and LIV. Design of vaccine based on subtype-specific strains may theoretically increase the vaccination efficient (for details, see Bukin et al. (2017)). However, this hypothesis ought to be tested in animals in the future.

## Notes

https://figshare.com/articles/dataset/Supplemental_tables/13619396

https://figshare.com/articles/dataset/BEAST_files/13614080

https://figshare.com/articles/dataset/TBFV_complete_genome_amino_acid_alignment/13613927

https://figshare.com/articles/figure/Supplemental_figures/13613888

### Competing Interest Statement

The authors have declared no competing interest.

## References

Adelshin, R.V., Melnikova, O.V., Karan, L.S., Andaev, E.I., Balakhonov, S.V., 2015. Complete genome sequences of four European subtype strains of tick-borne encephalitis virus from Eastern Siberia, Russia. Genome Announc 3. https://doi.org/10.1128/genomeA.00609-15.

Adelshin, R.V., Sidorova, E.A., Bondaryuk, A.N., Trukhina, A., Sherbakov, D.Y., White III, R.A., Andaev, E.I., Balakhonov, S.V., 2019. “886-84-like” tick-borne encephalitis virus strains: Intraspecific status elucidated by comparative genomics. Ticks Tick Borne Dis. 10, 1168–1172. https://doi.org/10.1016/j.ttbdis.2019.06.006.

Beasley, W.C., Suderman, T., Holbrook, R., Barrett, D.T., 2001. Nucleotide sequencing and serological evidence that the recently recognized deer tick virus is a genotype of Powassan virus. Virus Res. 79, 81–89. https://doi.org/10.1016/s0168-1702(01)00330-6.

Bogovic, P., Strle, F., 2015. Tick-borne encephalitis: A review of epidemiology, clinical characteristics, and management. World J. Clin. Cases 3, 430–441. https://doi.org/10.12998/wjcc.v3.i5.430.

Bukin, Y.S., Dzhioev, Y.P., Tkachev, S.E., Kozlova, I.V., Paramonov, A.I., Ruzek, D., Qu, Z., Zlobin, V.I., 2017. A comparative analysis on the physicochemical properties of tick-borne encephalitis virus envelope protein residues that affect its antigenic properties. Virus Res. 238, 124–132. https://doi.org/10.1016/j.virusres.2017.06.006.

Charrel, R.N., Zaki, A.M., Attoui, H., Fakeeh, M., Billoir, F., Yousef, A.I., de Chesse, R., De Micco, P., Gould, E.A., de Lamballerie, X., 2001. Complete coding sequence of the Alkhurma virus, a tick-borne flavivirus causing severe hemorrhagic fever in humans in Saudi Arabia. Biochem. Biophys. Res. Commun. 287, 455–461. https://doi.org/10.1006/bbrc.2001.5610.

Dai, X., Shang, G., Lu, S., Yang, J., Xu, J., 2018. A new subtype of eastern tick-borne encephalitis virus discovered in Qinghai-Tibet Plateau, China. Emerg. Microbes Infect. 7, 74. https://doi.org/10.1038/s41426-018-0081-6.

Demina, T.V., Dzhioev, Y.P., Verkhozina, M.M., Kozlova, I.V., Tkachev, S.E., Plyusnin, A., Doroshchenko, E.K., Lisak, O.V., Zlobin, V.I., 2010. Genotyping and characterization of the geographical distribution of tick-borne encephalitis virus variants with a set of molecular probes. J Med Virol 82, 965–976. https://doi.org/10.1002/jmv.21765.

Demina, T.V., Tkachev, S.E., Kozlova, I.V., Doroshchenko, E.K., Lisak, O.V., Suntsova, O.V., Verkhozina, M.M., Dzhioev, Y.P., Paramonov, A.I., Tikunov, A.Y., et al., 2017. Comparative analysis of complete genome sequences of European subtype tick-borne encephalitis virus strains isolated from Ixodes persulcatus ticks, long-tailed ground squirrel (Spermophilus undulatus), and human blood in the Asian part of Russia. Ticks Tick Borne Dis 8, 547–553. https://doi.org/10.1016/j.ttbdis.2017.03.002.

Dobler, G., Erber, W., Schmitt, H.J., 2017. TBE - The Book. Global Health Press, Singapore.

Fares, W., Dachraoui, K., Cherni, S., Barhoumi, W., Slimane, T.B., Younsi, H., Zhioua, E., 2020. Tick-borne encephalitis virus in Ixodes ricinus (Acari: Ixodidae) ticks, Tunisia. Ticks Tick Borne Dis. 12, 101606. https://doi.org/10.1016/j.ttbdis.2020.101606.

Fujisawa, T., Barraclough, T.G., 2013. Delimiting species using single-locus data and the Generalized Mixed Yule Coalescent approach: a revised method and evaluation on simulated data sets. Syst Biol 62, 707–724. https://doi.org/10.1093/sysbio/syt033.

Gilbert, L., 2016. Louping ill virus in the UK: a review of the hosts, transmission and ecological consequences of control. Exp. Appl. Acarol. 68, 363–374. https://doi.org/10.1007/s10493-015-9952-x.

Gilbert, L., Jones, L.D., Hudson, P.J., Gould, E.A., Reid, H.W., 2000. Role of small mammals in the persistence of Louping-ill virus: field survey and tick cofeeding studies. Med. Vet. Entomol. 14, 277–282. https://doi.org/10.1046/j.1365-2915.2000.00236.x.

Grard, G., Moureau, G., Charrel, R.N., Lemasson, J.J., Gonzalez, J.P., Gallian, P., Gritsun, T.S., Holmes, E.C., Gould, E.A., de Lamballerie, X., 2007. Genetic characterization of tick-borne flaviviruses: new insights into evolution, pathogenetic determinants and taxonomy. Virology 361, 80–92. https://doi.org/10.1016/j.virol.2006.09.015.

Gritsun, T.S., Lashkevich, V.A., Gould, E.A., 2003. Tick-borne encephalitis. Antivir. Res. 57, 129–146. https://doi.org/10.1016/s0166-3542(02)00206-1.

Heinze, D.M., Gould, E.A., Forrester, N.L., 2012. Revisiting the clinal concept of evolution and dispersal for the tick-borne flaviviruses by using phylogenetic and biogeographic analyses. J. Virol. 86, 8663–8671. https://doi.org/10.1128/JVI.01013-12.

Hoang, D.T., Chernomor, O., von Haeseler, A., Minh, B.Q., Vinh, L.S., 2018. UFBoot2: Improving the Ultrafast Bootstrap Approximation. Mol. Biol. Evol. 35, 518–522. https://doi.org/10.1093/molbev/msx281.

Holding, M., Dowall, S.D., Medlock, J.M., Carter, D.P., Pullan, S.T., Lewis, J., Vipond, R., Rocchi, M.S., Baylis, M., Hewson, R., 2020. Tick-borne encephalitis virus, United Kingdom. Emerg. Infect. Dis. 26, 90–96. https://doi.org/10.3201/eid2601.191085.

Hubálek, Z., Pow, I., Reid, H.W., Hussain, M.H., 1995. Antigenic similarity of central European encephalitis and louping-ill viruses. Acta Virologica 39, 251–256, (eng).

Hudson, R.R., Slatkin, M., Maddison, W.P., 1992. Estimation of levels of gene flow from DNA sequence data. Genetics 132, 583–589.

Jeffries, C.L., Mansfield, K.L., Phipps, L.P., Wakeley, P.R., Mearns, R., Schock, A., Bell, S., Breed, A.C., Fooks, A.R., Johnson, N., 2014. Louping ill virus: an endemic tick-borne disease of Great Britain. J. Gen. Virol. 95, 1005–1014. https://doi.org/10.1099/vir.0.062356-0.

Kaiser, R., 1999. The clinical and epidemiological profile of tick-borne encephalitis in southern Germany 1994-98: a prospective study of 656 patients. Brain 122 (Pt 11), 2067–2078. https://doi.org/10.1093/brain/122.11.2067.

Kalyaanamoorthy, S., Minh, B.Q., Wong, T.K.F., von Haeseler, A., Jermiin, L.S., 2017. ModelFinder: fast model selection for accurate phylogenetic estimates. Nat. Methods 14, 587–589. https://doi.org/10.1038/nmeth.4285.

Katoh, K., Rozewicki, J., Yamada, K.D., 2017. MAFFT online service: multiple sequence alignment, interactive sequence choice and visualization. Brief Bioinform 20, 1160–1166. https://doi.org/10.1093/bib/bbx108.

Kovalev, S.Y., Mukhacheva, T.A., 2017. Reconsidering the classification of tick-borne encephalitis virus within the Siberian subtype gives new insights into its evolutionary history. Infect., Genet. Evol. 55, 159–165. https://doi.org/10.1016/j.meegid.2017.09.014.

Kozlova, I.V., Demina, T.V., Tkachev, S.E., Doroshchenko, E.K., Lisak, O.V., Verkhozina, M.M., Karan, L.S., Dzhioev, Y.P., Paramonov, A.I., Suntsova, O.V., et al., 2018. Characteristics of the Baikal Subtype of Tick-Borne Encephalitis Virus Circulating in Eastern Siberia. Acta Biomedica Scientifica (East Siberian Biomedical Journal) 3, 53–60. https://doi.org/10.29413/abs.2018-3.4.9.

Kuraku, S., Zmasek, C.M., Nishimura, O., Katoh, K., 2013. aLeaves facilitates on-demand exploration of metazoan gene family trees on MAFFT sequence alignment server with enhanced interactivity. Nucleic Acids Res 41, W22–28. https://doi.org/10.1093/nar/gkt389.

Larsson, A., 2014. AliView: a fast and lightweight alignment viewer and editor for large data sets. Bioinformatics 30, 3276–3278. https://doi.org/10.1093/bioinformatics/btu531.

Leonova, G.N., Kondratov, I.G., Maystrovskaya, O.S., Takashima, I., Belikov, S.I., 2015. Louping ill virus (LIV) in the Far East. Arch. Virol. 160, 663–673. https://doi.org/10.1007/s00705-014-2310-1.

Mansfield, K.L., Johnson, N., Phipps, L.P., Stephenson, J.R., Fooks, A.R., Solomon, T., 2009. Tick-borne encephalitis virus - a review of an emerging zoonosis. J Gen Virol 90, 1781–1794. https://doi.org/10.1099/vir.0.011437-0.

Miller, M.A., Schwartz, T., Pickett, B.E., 2010. Creating the CIPRES Science Gateway for inference of large phylogenetic trees. Gateway Computing Environments Workshop (GCE), 2010. IEEE, pp. 1–8.

Moureau, G., Cook, S., Lemey, P., Nougairede, A., Forrester, N.L., Khasnatinov, M., Charrel, R.N., Firth, A.E., Gould, E.A., de Lamballerie, X., 2015. New insights into flavivirus evolution, taxonomy and biogeographic history, extended by analysis of canonical and alternative coding sequences. PLoS One 10, e0117849. https://doi.org/10.1371/journal.pone.0117849.

Nguyen, L.T., Schmidt, H.A., von Haeseler, A., Minh, B.Q., 2015. IQ-TREE: a fast and effective stochastic algorithm for estimating maximum-likelihood phylogenies. Mol. Biol. Evol. 32, 268–274. https://doi.org/10.1093/molbev/msu300.

Pickett, B.E., Sadat, E.L., Zhang, Y., Noronha, J.M., Squires, R.B., Hunt, V., Liu, M., Kumar, S., Zaremba, S., Gu, Z., et al., 2012. ViPR: an open bioinformatics database and analysis resource for virology research. Nucleic. Acids Res. 40, D593–598. https://doi.org/10.1093/nar/gkr859.

Puillandre, N., Lambert, A., Brouillet, S., Achaz, G., 2012. ABGD, Automatic Barcode Gap Discovery for primary species delimitation. Mol. Ecol. 21, 1864–1877. https://doi.org/10.1111/j.1365-294X.2011.05239.x.

Rambaut, A., Drummond, A.J., Dong, X., Baele, G., Suchard, M.A., 2018. Posterior summarization in Bayesian phylogenetics using tracer 1.7. Syst. Biol. 67, 901–904. https://doi.org/10.1093/sysbio/syy032.

Rey, F.A., Heinz, F.X., Mandl, C., Kunz, C., Harrison, S.C., 1995. The envelope glycoprotein from tick-borne encephalitis virus at 2 A resolution. Nature 375, 291–298. https://doi.org/10.1038/375291a0.

Shi, J., Hu, Z., Deng, F., Shen, S., 2018. Tick-Borne Viruses. Virol. Sin. 33, 21–43. https://doi.org/10.1007/s12250-018-0019-0.

Suchard, M.A., Lemey, P., Baele, G., Ayres, D.L., Drummond, A.J., Rambaut, A., 2018. Bayesian phylogenetic and phylodynamic data integration using BEAST 1.10. Virus Evol. 4, vey016. https://doi.org/10.1093/ve/vey016.

Tonteri, E., Kipar, A., Voutilainen, L., Vene, S., Vaheri, A., Vapalahti, O., Lundkvist, A., 2013. The three subtypes of tick-borne encephalitis virus induce encephalitis in a natural host, the bank vole (Myodes glareolus). PLoS One 8, e81214. https://doi.org/10.1371/journal.pone.0081214.

Uzcátegui, N.Y., Sironen, T., Golovljova, I., Jääskeläinen, A.E., Välimaa, H., Lundkvist, Å., Plyusnin, A., Vaheri, A., Vapalahti, O., 2012. Rate of evolution and molecular epidemiology of tick-borne encephalitis virus in Europe, including two isolations from the same focus 44 years apart. J. Gen. Virol. 93, 786–796. https://doi.org/10.1099/vir.0.035766-0.

Votiakov, V.I., Protas, I.I., Zhdanov, V.M., 1978. Western Tick-Borne Encephalitis. Belarus, Minsk.

Votiakov, V.I., Zlobin, V.I., Mishaeva, N.P., 2002. Tick-borne encephalitis of Eurasia (ecology, molecular epidemiology, nosology, evolution). Nauka, Novosibirsk.

Williams, H., Thorburn, H., 1962. Serum antibodies to louping-ill virus. Scott. Med. J. 7, 353–355. https://doi.org/10.1177/003693306200700803.

Zhang, J., Kapli, P., Pavlidis, P., Stamatakis, A., 2013. A general species delimitation method with applications to phylogenetic placements. Bioinformatics 29, 2869–2876. https://doi.org/10.1093/bioinformatics/btt499.

