## Supplementary figures and images for "Delimitation of the Tick-Borne Flaviviruses. Resolving the Tick-Borne Encephalitis and Louping-Ill Virus Paraphyletic Taxa"

### supplemental fig 1

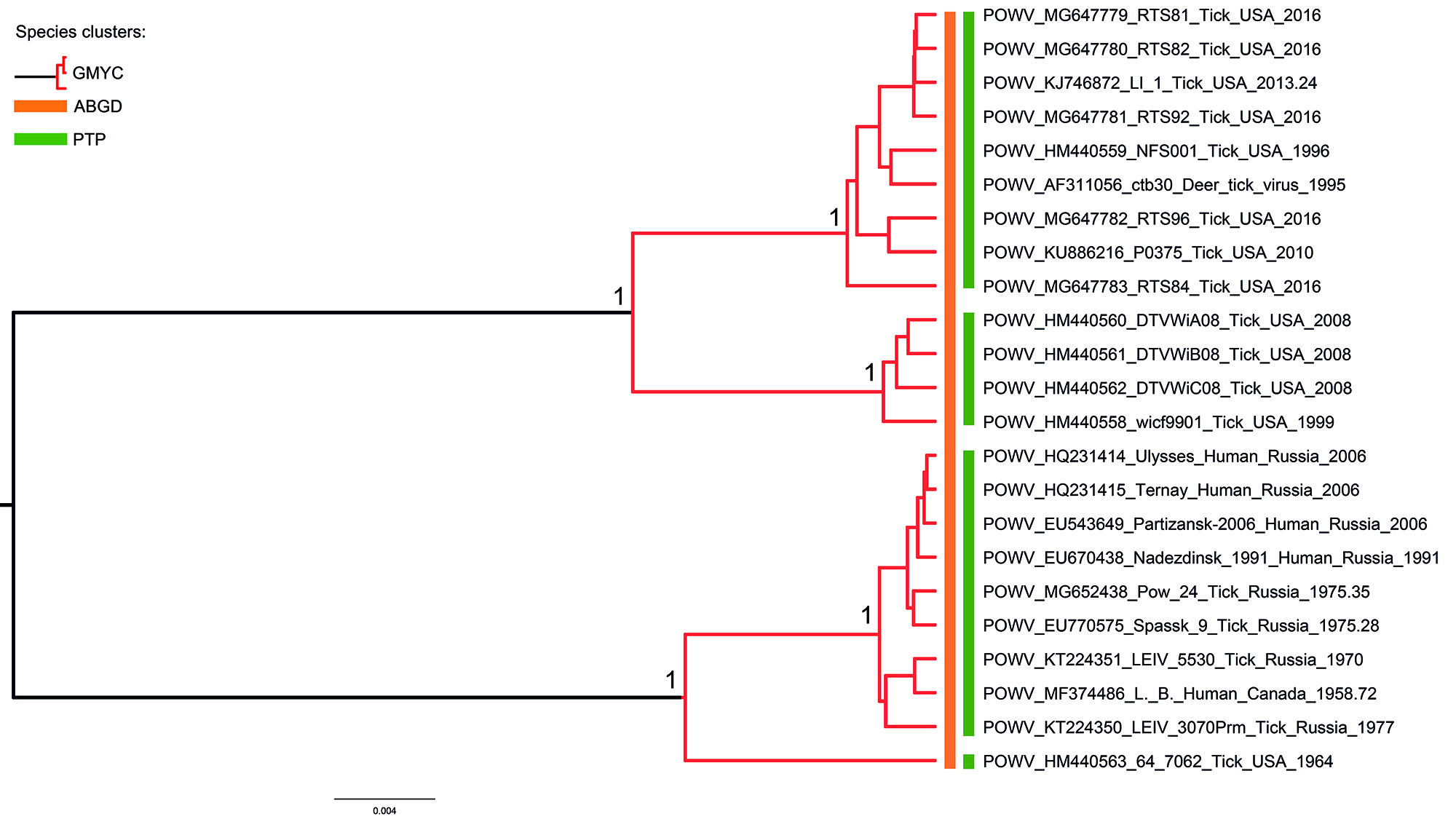

### supplemental fig 2

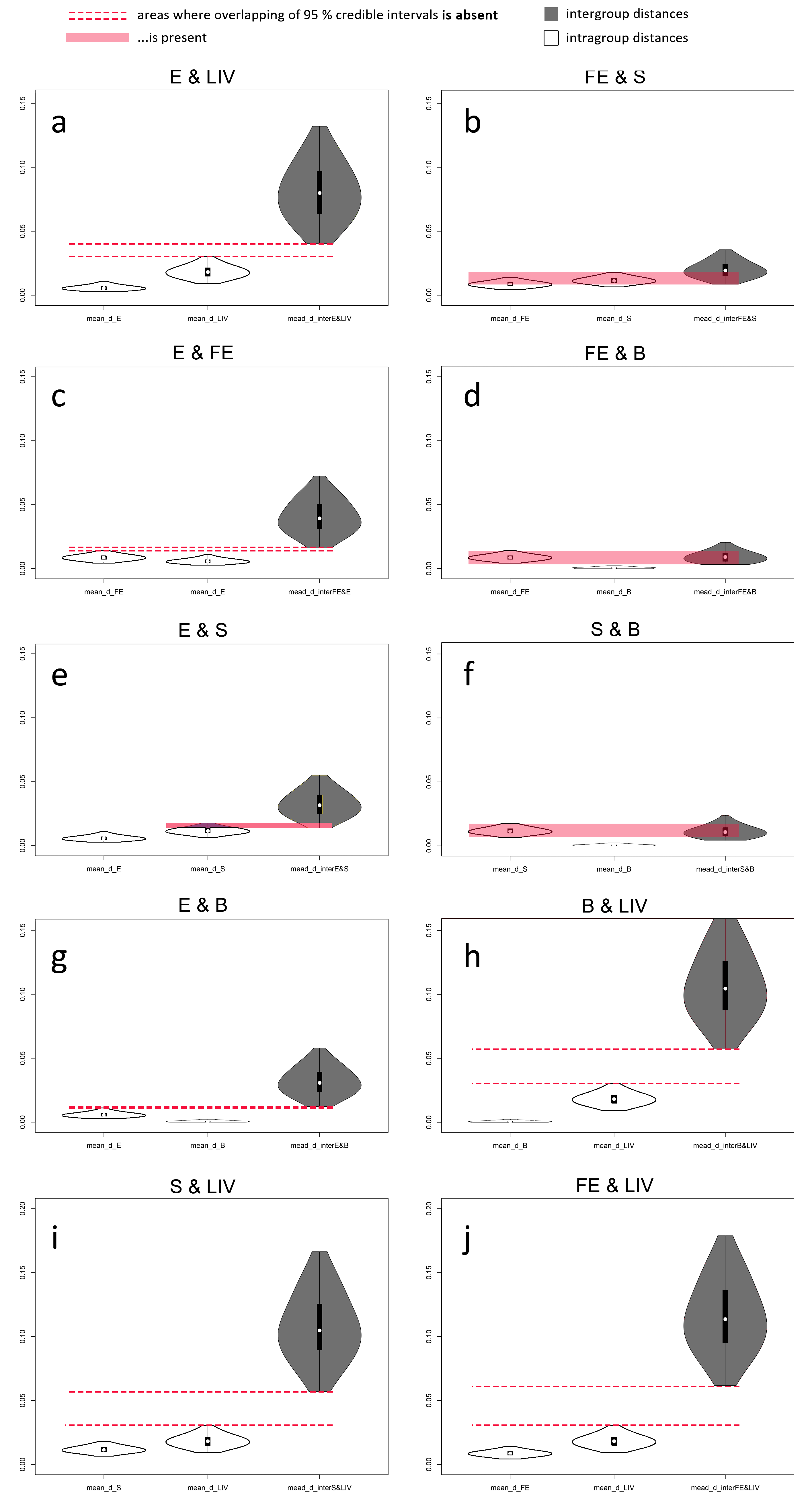

### supplemental fig 3

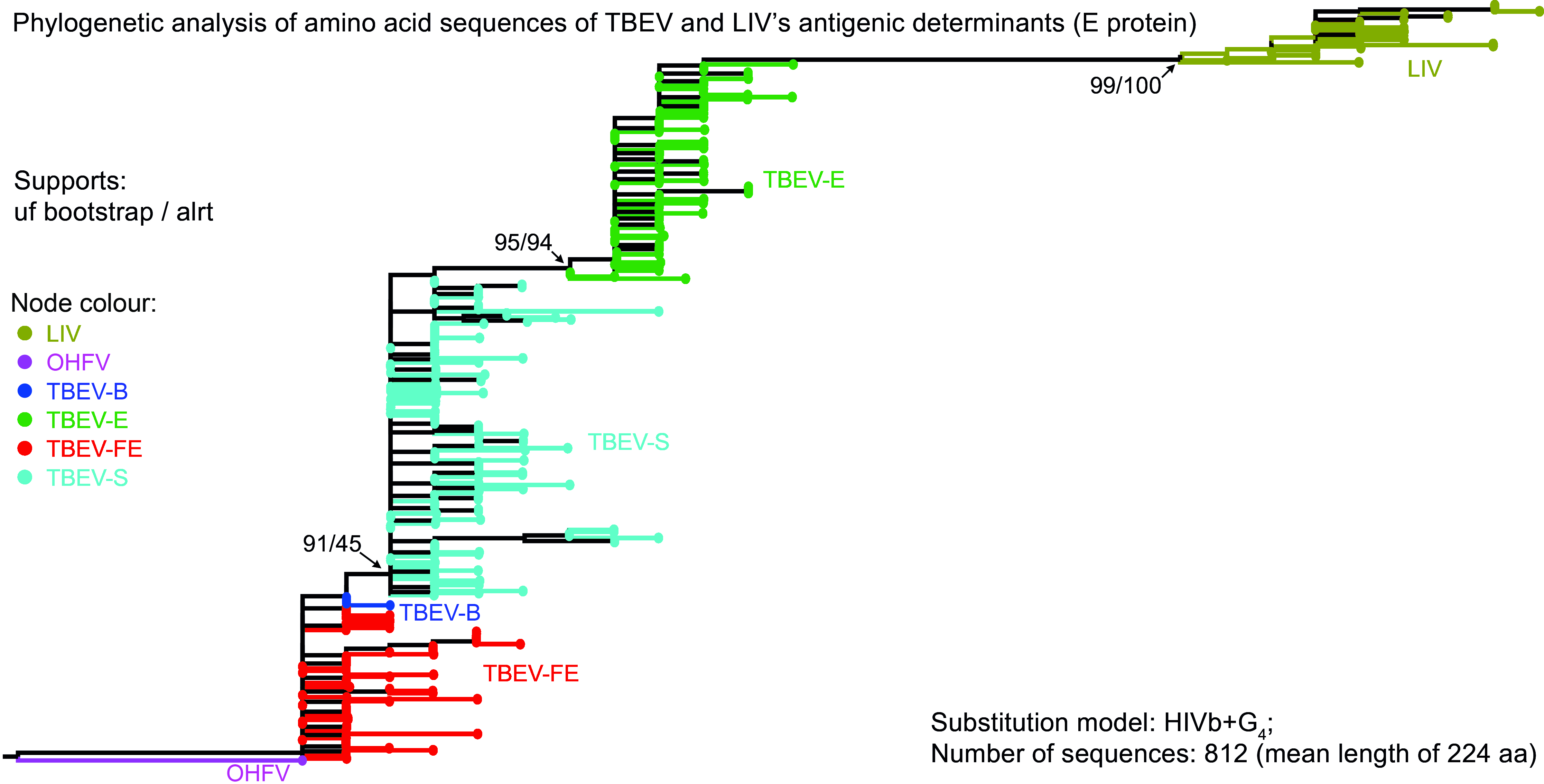
